# RobPicker: A Meta Learning Framework for Robust Identification of Macromolecules in Cryo-Electron Tomograms

**DOI:** 10.1101/2025.09.16.676650

**Authors:** Ramtin Hosseini, Youwei Liang, Digvijay Singh, Hamidreza Rahmani, Sang Choe, Joan Lee, Min Xu, Eran Segal, James Zou, James Williamson, Danielle A. Grotjahn, Elizabeth Villa, Pengtao Xie

## Abstract

Accurate particle picking of macromolecules of all kinds, shapes, and sizes in cryogenic electron tomograms (cryo-ET) is critical for understanding the molecular architecture of biological systems in their native state. Current deep learning methods have shown potential in identifying macromolecules from tomograms, but they are vulnerable to issues of noise in the training dataset obtained through human or automatic labeling and imbalanced distribution of macromolecule species. To address these limitations, we developed RobPicker, a meta-learning framework that effectively mitigates these issues by automatically learning deep neural networks to correct label errors and give greater emphasis to underrepresented macromolecule species. In evaluations across diverse cryo-ET datasets with noisy labels and imbalanced species distributions, RobPicker substantially outperforms state-of-the-art methods, particularly in identifying small and rare macromolecules. The efficiency and robustness of RobPicker can also be used for rapid fine-tuning of the tilt-series alignment, leading to improved tomogram reconstruction and enabling high-resolution cellular structural biology analysis.

## Introduction

Cryo-electron tomography (cryo-ET) has revolutionized structural biology by enabling the high-resolution visualization and structure determination of macromolecules in their near-native states within the complex environment of cells in situ [1–8]. Accurately identifying and locating macromolecules in cryo-ET is crucial for understanding their functions, interactions, and dynamics within the cellular context [9]. With the advance of instrumentation [3, 10], data processing [11, 12], and automation [13, 14], the volume of in-cell cryo-ET data is expanding rapidly in public databases like EMPIAR [15]. By training neural networks on annotated tomograms, recent deep learning methods have shown promise in automatically picking macromolecule complexes (referred to as particles) from cryo-ET data [16–23]. In particular, macromolecule segmentation networks, in particular 3D U-Net [24], have been successfully applied to segment and pick particles such as ribosomes [20, 21]. Yet, these networks often struggle with robustness, particularly facing two key challenges: noise in the training labels brought in by human or automatic annotations and imbalanced distribution of different species [22, 25].

The low signal-to-noise ratio (SNR) of cryo-ET raw data, combined with incomplete angular sampling due to the physical constraints of the microscope (i.e., the ‘missing wedge’ artifact [26]) presents significant challenges in detecting and classifying particles within the crowded cellular milieu, especially those of smaller size [27]. Recent cryo-ET denoising methods [28–30] have improved the SNR of tomograms but they still cannot remove the noise [31]. The noisy tomograms also complicate the annotation of particles, which is necessary for training supervised deep learning models for particle picking. Typical particle annotation methods include manual, semi-automated, and automated annotations. In manual annotation, human experts manually browse the tomogram and mark the locations of particles. While manual annotation can be accurate for large particles, it is extremely labor intensive [32]. Therefore, semi-automated and automated annotation pipelines have been developed to accelerate the annotation process, using template matching [33] and deep learning models [21, 32, 34]. However, template matching often results in false positives [35], and supervised deep learning models struggle with identifying small particles, especially when the training data is scarce [36]. In summary, particle annotation in cryo-ET is inherently difficult (especially for small particles) and time-consuming, while automated annotations often lead to inconsistent or erroneous labels (referred to as noisy labels). When the noisy labels are used for deep learning model training, the resulting models cannot accurately identify particles when applied to new tomograms (Fig. 1a).

**Fig. 1:**
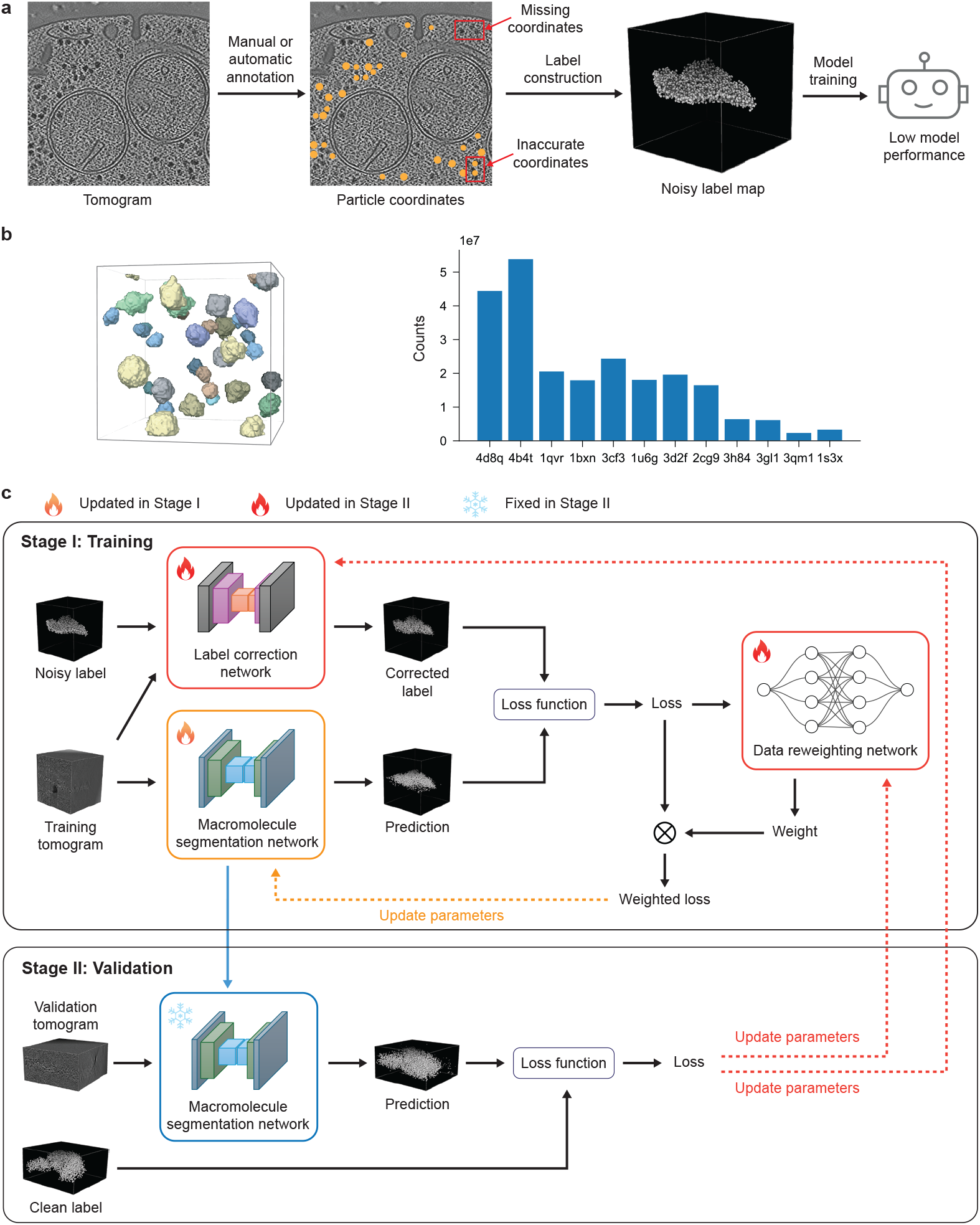
Noisy labels and species imbalance are two common issues that decrease the accuracy of macromolecular segmentation models for deep learning (DL)-based particle picking. To address these issues, RobPicker’s bi-level optimization involving data reweighting and labeling correction enables the robust segmentation of macromolecules. **a**, An illustration of noisy labels on a tomographic slice. Manual or automatic annotation often results in missing and inaccurate coordinates for target particles (yellow dots are annotations), which further leads to noisy label maps in typical cryo-ET training data. The resulting DL model does not generalize well to particle picking in new tomograms. **b**, The left panel shows the label map of a typical subtomogram in the SHREC 2019 dataset. Different colors denote different macromolecule species. Larger particles occupy more voxels in the label. The right panel shows the number of voxels for each species in the SHREC 2019 dataset, which shows the distribution is highly imbalanced. Models are less trained on the labels of the smaller macromolecules, resulting in lower model performance on smaller macromolecules. **c**, RobPicker comprises three components: a macromolecule segmentation network, a data reweighting network, and a label correction network, all integrated into a two-stage end-to-end learning process. In Stage I, given a tomogram and its noisy macromolecule labels, the segmentation network detects macromolecules, and the label correction network refines the labels. The prediction loss, computed between the detection results and the corrected labels, is then passed to the data reweighting network, which generates a weight that scales the prediction loss. The segmentation network is trained by minimizing the weighted losses across all training examples. In Stage II, the segmentation network, trained during Stage I, is evaluated on the validation tomograms with clean labels. The validation loss, which depends on both the data reweighting and label correction networks, is minimized to optimize these two networks.

Training macromolecule segmentation networks is further impeded by the significant imbalance of the occurrence of different macromolecular species within the cellular environment [37]. Larger, more abundant species occupy more voxels in the tomograms and tend to have a clearer boundary against the background, while smaller or less abundant species—though often playing vital roles in cellular processes—are less represented and occupy fewer voxels. As training macromolecule segmentation networks mostly relies on voxel-wise loss functions—such as Dice loss [38, 39]—to update the model parameters, larger and more abundant species have a larger influence on the model update [40]. This imbalance leads to a skewed performance where models are proficient at identifying large, well-delineated species but struggle with picking smaller, rarer ones (Fig. 1b). The inability to consistently pick small particles in cryo-ET data has hindered our understanding of smaller macromolecules and their critical roles in biological systems.

To address these limitations, we introduce RobPicker, a novel meta-learning [41] framework that improves the robustness of macromolecule segmentation networks for cryo-ET. Unlike traditional supervised deep learning methods, which rely heavily on vast quantities of well-labeled data, RobPicker is designed to overcome challenges posed by labeling noise and imbalanced species distribution. It achieves this by automatically learning a label correction network that rectifies labeling errors and a data reweighting network that gives greater emphasis to underrepresented macromolecule species, which is guided by evaluating the performance of the macromolecule segmentation network on a validation set with cleaner labels and a more balanced species distribution. These corrected and reweighted data are then used to learn a more robust macromolecule segmentation network (Fig. 1c). Through comprehensive evaluations of both experimental cryo-ET data and benchmarks, RobPicker demonstrates significantly improved robustness in identifying particle species in various cell types even when the labels are noisy and the species distribution is imbalanced. Moreover, we demonstrated the effectiveness of RobPicker by utilizing it to pick a set of initial ribosome particles efficiently and robustly, which were further used for quick multiple-particle refinement [42], enabling a fast fine-tuning of the alignment of tilt-series.

## Results

### RobPicker overview

RobPicker is composed of three deep neural networks: a macromolecule segmentation network, a label correction network, and a data reweighting network. The macromolecule segmentation network takes a cryo-electron tomogram as input and outputs the detection result in the form of a segmentation map, which indicates the likelihood of macromolecule species at each voxel in the tomogram. The segmentation map is used to derive the class of species and the center of particles in post processing (Methods). While we employed a 3D U-Net [24] as the macromolecule segmentation network—following prior deep learning pickers [20], our method can benefit from using more advanced segmentation network architectures.

The label correction network is designed to correct noisy labels in the training data. It is a small 3D U-Net that takes a tomogram and a noisy particle segmentation map as inputs and produces a corrected label (Fig. 1c and Methods). The parameters of the label correction network are updated to minimize the segmentation loss function on a separate, smaller dataset (called validation set) with cleaner—compared to the training set—labels [43] (Stage II in Fig. 1c). Intuitively, a bad label correction network will result in corrupted supervision signal for the segmentation network to learn in the training stage (Stage I in Fig. 1c), which will result in low segmentation performance in the validation stage—thus a high validation loss—since the validation labels are cleaner. Therefore, minimizing the validation loss can update the label correction network towards the direction such that it outputs better supervision signal, i.e., cleaner labels. The experiment on the SHREC 2019 cryo-ET dataset [44] revealed that the trained label correction network recovered missing labels of ground truth particles. An exemplar corrected label map is shown in Fig. 2a, underscoring the effectiveness of the label correction mechanism in RobPicker.

**Fig. 2:**
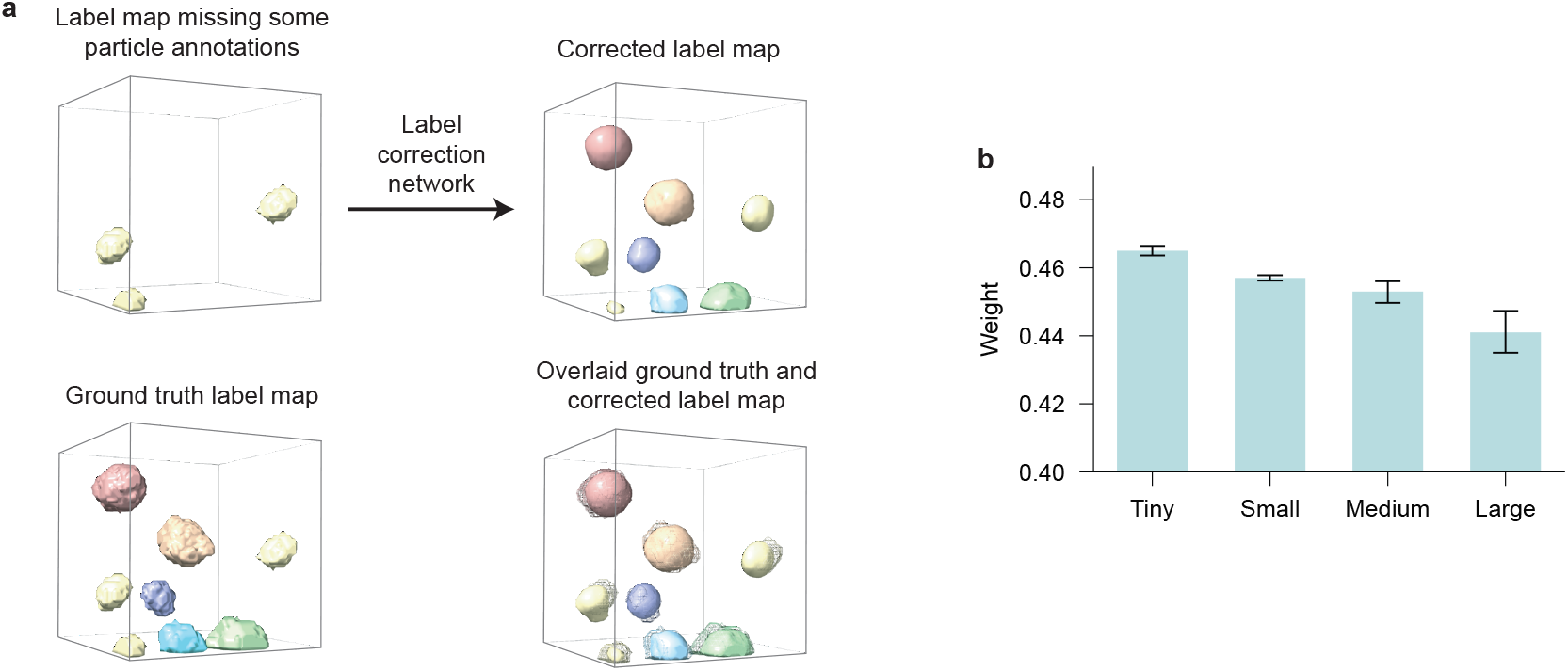
RobPicker’s label correction network and data reweighting network effectively correct missing labels and assign higher weights to minority species in the SHREC 2019 dataset. **a**, Based on a tomogram and a noisy label map with some missing particle annotations, RobPicker’s label correction network predicted a new label map that recovered the missing particle labels. The overlaid ground truth (shown in mesh) and corrected label (shown in solid objects) show high alignment. Different colors denote different species. **b**, RobPicker’s data reweighting network assigned larger weights—the mean and standard deviation are shown—to smaller macromolecule species, accounting for the data imbalance. The x-axis denotes particle sizes.

In parallel to the label correction network, the data reweighting network [45] predicts a weight for each tomogram to increase the influence of smaller or less abundant particle species on the update of the segmentation network. Specifically, the reweighting network takes as input the prediction loss of a tomogram and outputs a scalar weight between 0 and 1, which is multiplied by the prediction loss so that the weight controls the influence—higher weights lead to higher influence—of the tomogram on model parameter update (Stage I in Fig. 1c). The reweighting network is implemented as a multi-layer perceptron (MLP) with one hidden layer. Similar to the update of the label correction network, the update of the reweighting network is also guided by minimizing the validation loss on the validation set where balanced data sampling is used to increase the frequency of less abundant species (Methods). The experiment on the SHREC 2019 cryo-ET dataset [44] revealed that the trained data reweighting network assigned larger weights to tomograms whose major particles belong to the smaller particle species (Fig. 2b), confirming the effectiveness of the reweighting mechanism in RobPicker.

### RobPicker tolerates noise in cryo-ET particle picking training data

Typical cryo-ET particle picking datasets contain annotations of particle locations in the form of their 3D coordinates. Following DeepFinder’s data preprocessing approach, we applied two methods—a sphere-based and a shape-based—to derive label maps from the location annotations [20]. The shape-based method uses subtomogram averaging [13] to create a mask that mimics the shape of each particle species, and then the mask is placed at each particle location to represent the particles in the label map. The sphere-based method simply uses a sphere mask to represent each particle in the label map, which is more computationally efficient than the shape-based method but resulting in noisier labels. Following the widely used protocol for model training and evaluation [20], we divided our cryo-ET datasets into three subsets: a training set, a validation set, and a test set. We aim to experiment with using the sphere-based method to efficiently generate labels for the majority of the tomograms and examine if RobPicker can tolerate the noise brought in by the sphere-based labels. To this end, we used the sphere-based method to create the label maps for the tomograms in the training set (which has the majority of the tomograms), while we used the shape-based method to create the label maps for the remaining small amount of tomograms and put them in the validation and test sets. The data preprocessing is efficient since subtomogram averaging is only done for a small amount of tomograms.

We conducted experiments on four cryo-ET real datasets (denoted by D1–D4) comprising tomograms with annotations for ribosomes from different cell types. The first dataset (D1) contains annotations for cytosolic ribosomes (ct-ribos) and membrane-bound 80S ribosomes (mb-ribos) in *C. reinhardtii* cells [20] (detailed statistics can be found in Table 1). D2 was annotated for 80S ribosomes in *S. cerevisiae* (yeast) cells [46]. D3 contains annotations for 50S large subunits and fully assembled 70S ribosomes of *E. coli* cells. D4 also contains annotations for ribosomes in yeast cells. The details of data annotation can be found in the Methods section. We compared RobPicker against a state-of-the-art supervised deep learning method DeepFinder [20], which uses a 3D U-Net [24] for multi-class semantic segmentation to pick particles in tomograms. We used the same 3D U-Net as the segmentation network in RobPicker as DeepFinder to demonstrate that the robustness of RobPicker results from the training framework (Fig. 1c) instead of more advanced network architectures. Therefore, the major distinction between RobPicker and DeepFinder is that RobPicker leveraged a data reweighting network and a label correction network to reweight and correct the training data. To report model performance, we calculated the picking F1 scores as the evaluation metric as in DeepFinder. In the evaluation, a particle prediction is considered as a true positive only when the prediction and the ground truth have sufficient overlap (Methods). The F1 score ranges from 0 (lowest performance) to 1 (highest performance). The experiments were repeated three times, and the mean and standard deviation are reported in Fig. 3a–f.

**Table 1.** Dataset statistics. We report the number of tomograms, and number of annotated particles in the training, validation, and test sets.

| Dataset | Object | Training | Validation | Test |
| --- | --- | --- | --- | --- |
| D1 | tomogram | 48 | 1 | 8 |
|  | ct-ribos | 6,687 | 254 | 2,594 |
|  | mt-ribos | 6,834 | 222 | 1,736 |
| D2 | tomogram | 4 | 1 | 1 |
|  | ribosome | 2,986 | 707 | 843 |
| D3 | tomogram | 26 | 1 | 1 |
|  | 50S | 6,013 | 443 | 238 |
|  | 70S | 6,318 | 512 | 285 |
| D4 | tomogram | 3 | 1 | 1 |
|  | ribosome | 6,467 | 1,238 | 1,496 |

**Fig. 3:**
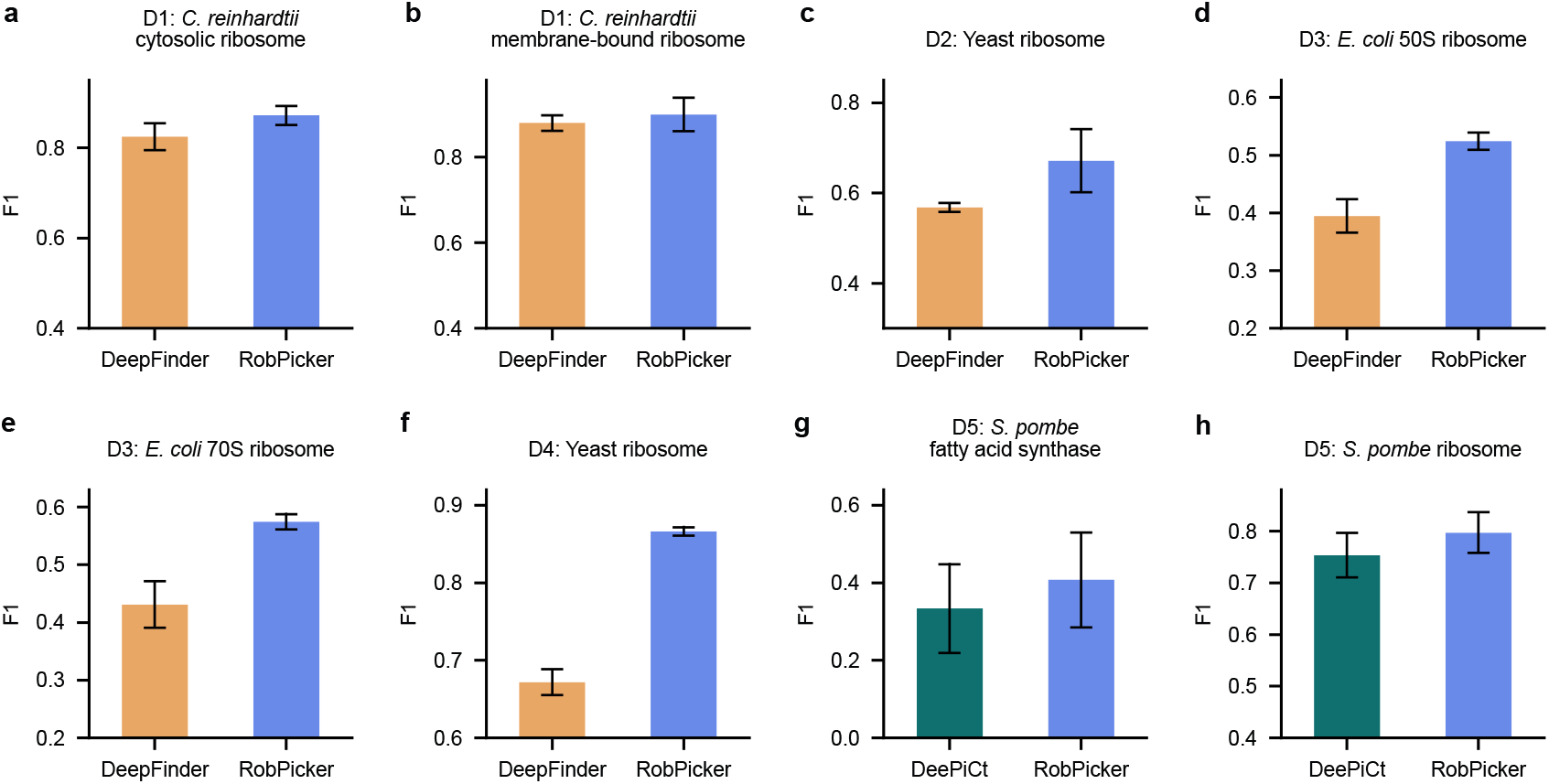
RobPicker achieved higher mean picking F1-scores than DeepFinder and DeePiCt in various cryo-ET real datasets. The mean and standard deviation are shown.

For identification of ct-ribos in D1, RobPicker achieved a mean F1 score of 0.87, outperforming DeepFinder’s mean F1 score of 0.82 (*P* = 0.098). For mb-ribos in D1, RobPicker obtained a mean F1 score of 0.9, surpassing DeepFinder’s F1 score of 0.88 (*P* = 0.48). For yeast ribosome identification in D2, RobPicker achieved a mean F1 score of 0.67, outperforming DeepFinder’s F1 score of 0.57 by 17.5% (*P* = 0.12). In D3, RobPicker achieved mean F1 scores of 0.52 and 0.57 in the identification of *E. coli* 50S and 70S ribosomes, respectively. RobPicker substantially outperformed DeepFinder’s mean F1 scores of 0.39 (*P* = 0.0063) and 0.43 (*P* = 0.018), marking 33.3% and 32.6% relative improvement on picking 50S and 70S ribosomes, respectively. For yeast ribosome identification in D4, RobPicker achieved a mean F1 score of 0.87, considerably exceeding DeepFinder’s F1 score of 0.67 (*P* = 0.0010), which represents a 29.9% relative improvement. The performance comparison on the four real cryo-ET datasets show that RobPicker demonstrated substantially enhanced performance over DeepFinder in picking particles across diverse datasets. The experiments underscore that RobPicker tolerated the label noise in the training set robustly, enabling efficient data preprocessing for the training data, as only a smaller amount of clean labels is needed to calculate the validation loss of RobPicker. In contrast, the supervised learning method DeepFinder [20] achieved lower performance when the training data contains noisy labels created from sphere-based particle masks. The robustness of RobPicker can be attributed to the label correction mechanism in RobPicker, where the sphere-based particle labels in the training set are rectified by the label correction network to better represent the particle shapes, thus resulting in a more accurate macromolecule segmentation model.

### RobPicker robustly picks small and rare macromolecular species

We experimented with a cryo-ET real dataset (denoted by D5) obtained from DeePiCt [21], which contains 10 tomograms acquired from wild-type *S. pombe* using a Volta potential phase plate (VPP). It contains annotations of 731 fatty acid synthases (FAS) and 25,311 ribosomes, and highquality label masks of the macromolecules were generated for model training and evaluation [21]. We trained DeePiCt and RobPicker to localize the *S. pombe* FAS and ribosomes. FAS segmentation is particularly challenging due to its sporadic presence in cells [21]. We followed the original work [21] to use five-fold cross-validation, splitting the tomograms into five equal-sized subsets. In each fold, one subset was reserved for testing, while the remaining four were used for training. For RobPicker, the training set (8 tomograms) was further divided into a new training set and a validation set in a 7:1 ratio, used for Stage I and Stage II optimizations, respectively (Fig. 1c). We adhered to the same pre-processing and post-processing configurations as DeePiCt to ensure consistency. The mean and standard deviation of the picking F1 scores are shown in Fig. 3g-h.

RobPicker achieved a mean F1 score of 0.41 for FAS picking, marking a 24.2% relative improvement over DeeP-iCt’s mean F1 score of 0.33 (*P* = 0.181). For ribosome picking, RobPicker’s mean F1 score of 0.80 also surpasses DeePiCt’s mean F1 score of 0.75 (*P* = 0.029) with a 6.67% relative improvement. These results highlight RobPicker’s higher performance in particle picking compared to DeePiCt. Since only 731 FAS particles are present in the dataset, which is much fewer than the abundant ribosomes with a total of 25,311 particles, the species distribution is extremely imbalanced. Moreover, the FAS particles are much smaller than the ribosome particles. These factors make picking FAS much more challenging than picking ribosomes. Nevertheless, the relative improvement of the F1 scores of RobPicker over DeePiCt is much more significant on the FAS than on the ribosome (24.2% vs 6.67%), underscoring RobPicker’s robustness on picking smaller and less abundant particles. RobPicker’s robustness to imbalanced species distribution is attributed to the inclusion of the data reweighting network, which dynamically up-weight the minority species in training. This strategy is in contrast to traditional supervised learning methods like DeePiCt, which directly optimizes the segmentation loss function without reweighting the data samples, causing the segmentation network biases towards the major species in the segmentation.

### RobPicker enables efficient, supervision-free particle picking for macromolecular structure determination

The output of particle picking from cryo-ET data is often used as input for structure solving analysis, such as subtomogram averaging. We compared RobPicker against other particle picking methods in structure determination pipelines, which demonstrated RobPicker as a fast, first-round picking method without any supervision to get a high-quality 3D reconstruction (also referred to as 3D map or map).

Our cryo-ET data processing pipeline integrates RobPicker-based particle picking, subtomogram averaging, and multi-particle refinement [42] to efficiently achieve high-resolution reconstructions (Fig. 4). To demonstrate the pipeline’s effectiveness, we first trained RobPicker to pick ribosomes on just four yeast tomograms in D4, where RobPicker achieved a much higher F1 score than DeepFinder (0.87 vs 0.67) in the held-out yeast tomogram (Fig. 3f). As illustrated in Fig. 4a, RobPicker produced more accurate particle probability maps compared to DeepFinder, with significantly fewer false positives, reflecting its strong specificity. Furthermore, we applied RobPicker on another eight held-out tomograms of yeast, where the network inference was efficiently done in ∼15 minutes on an A100 GPU. We performed a quick 3D refinement followed by 3D classification on the picked particles in RELION [47], resulting in a good and representative yeast ribosome map (resolution ∼19 Å) shown in Fig. 4b, further underscoring RobPicker’s efficiency, accuracy, and specificity.

**Fig. 4:**
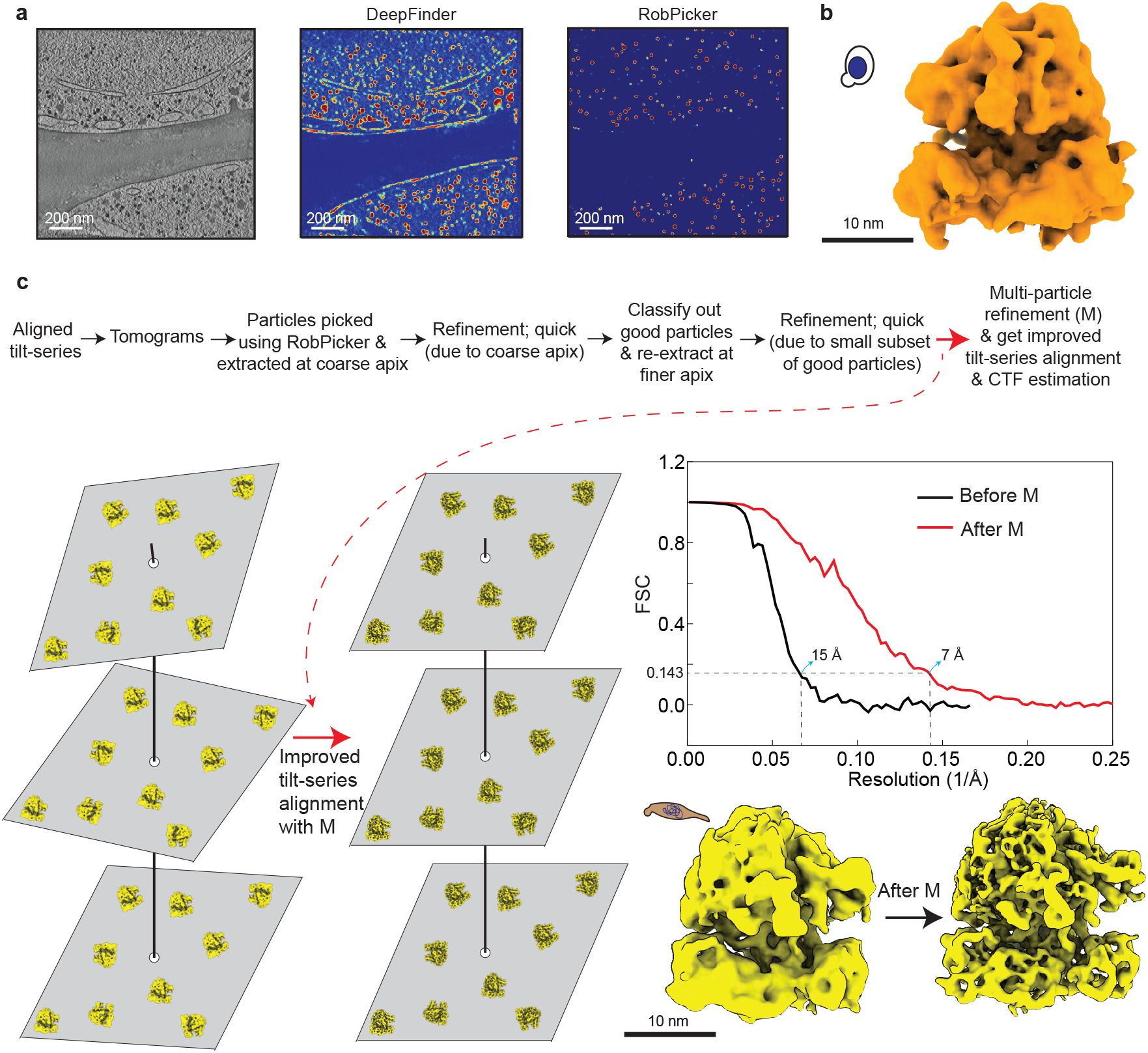
RobPicker’s accuracy and efficiency can be utilized for fast improvement of tilt-series alignment. **a**, Tomographic slice (1st column) of a representative tomogram from yeast, along with the heatmap showing the distribution of reference (ribosome) localization probabilities for DeepFinder and RobPicker. RobPicker achieved significantly higher specificity than DeepFinder. **b**, A de novo RobPicker model, which was trained on only 4 yeast tomograms, was used to pick ribosomes across other yeast tomograms. Quick refinement gave this map of the yeast ribosome, further underscoring RobPicker’s accuracy and specificity. **c**, The same RobPicker model for yeast ribosomes was used to pick ribosomes in in-cell bacterial tomograms at a highly coarse pixel size (apix). The picks produced a good structure of the bacterial ribosome, demonstrating the wide generalizability of RobPicker’s particle picking ability across different species and its ability to accurately predict on heavily down sampled (highly coarse apix) tomograms. Such picks were subjected to a multiple-particle refinement workflow shown in the top panel. All of this was performed quickly and led to a significant improvement in the resolution of the ribosome average. This process improved and fine-tuned the existing alignment of the tilt-series and its CTF estimation, showcasing how RobPicker can be used for routine and quick fine-tuning and improvement of the tilt-series alignments.

To evaluate RobPicker’s generalization capacity, the same RobPicker model trained on the four yeast tomograms was used to pick ribosomes in 14 in-cell tomograms at a highly coarse pixel size (apix) of another organism (bacterial; *M. pneumoniae* [42]). Subsequently, we aimed to create a high-resolution bacterial ribosome map using a pipeline subjected to Fig.4c. Specifically, we performed a 3D refinement with the picked particles in RELION [47], which was quick due to the coarse apix. The 3D classification resulted in a small subset of high-quality particles. These particles were re-extracted at finer apix for further refinement. To further improve and fine-tune tilt-series alignment and its resulting reconstruction quality, we used the multi-particle refinement tool M [42], which jointly optimizes tilt-series alignment and contrast transfer function (CTF) parameters across the full tomograms. Application of M to the refined bacterial ribosome particles resulted in a marked resolution improvement from ∼15 Å to 7 Å at the 0.143 Fourier shell correlation (FSC) criterion (Fig.4c). This whole process that led to a significant improvement in the tilt-series alignment and resolution of the ribosome average can be performed quickly within a few hours. This highlights the utility of RobPicker for fast, accurate particle detection and its synergy with modern refinement tools in enhancing tilt-series alignment and structural resolution.

We performed an experiment on another yeast tomogram dataset to compare traditional template matching methods and RobPicker in solving ribosome structures. The lack of supervision and manual curation required to clean up the picks is a major advantage of RobPicker. Such manual curation is a significant bottleneck for traditional templatematching methods like PyTom [48, 49] for particle picking. The manual curation is required in such template matching methods due to prevalence of false positives in the picked particles. But even this manual curation is ill-defined. Typically, a user-defined threshold (on quality score of template matching picks) has to be applied to pick high quality particles (presumably representing true positives). In contrast, RobPicker’s detection of ribosomes requires no manual input and can generalize well to other organisms without changing any hyper-parameters (as shown in the bacterial ribosome experiments above). After training RobPicker on five yeast tomograms in D2, we used another four heldout yeast tomograms as the test dataset for a comparison between PyTom and RobPicker. While RobPicker identified 4,092 particles in the four tomograms, the automatic thresholding in PyTom returned very few particles. This can be explained by the fact that there is no visible second peak in the histogram of the local cross-correlation (LCC) scores when we extract the top 10,000 candidates (Extended Data Fig. 1a). To compare the two particle-picking methods, we extracted the same number of particles as the RobPicker output from PyTom candidates (breakdown in Extended Data Fig. 1b). The comparison between the RobPicker picks and PyTom picks, shown in Fig. 5b–c, demonstrates the higher specificity of the RobPicker picks. We performed several rounds of refinement and classification in RELION [47] and further multi-particle refinement in M [42], for both particle sets to select a subset of particles that showed consistent high-resolution features during iterative refinement. This process resulted in an 11 Å reconstruction from the RobPicker-identified dataset, and a 12.5 Å reconstruction from the PyTom-identified dataset. Visual inspection of the reconstructions (Fig. 5d) shows a marked increase in detail in the map from RobPicker-identified picks compared to those identified in PyTom, emphasizing the improved ability of RobPicker to identify more particles that contribute to highresolution information in the resulting reconstruction. The ability to use RobPicker as a pre-trained model offers the potential of improving and speeding this process up even further for initial reconstruction of 80S ribosomes that can be used to improve our tomograms in packages like RELION and M, similar to the pipeline described in Fig. 4c.

**Fig. 5:**
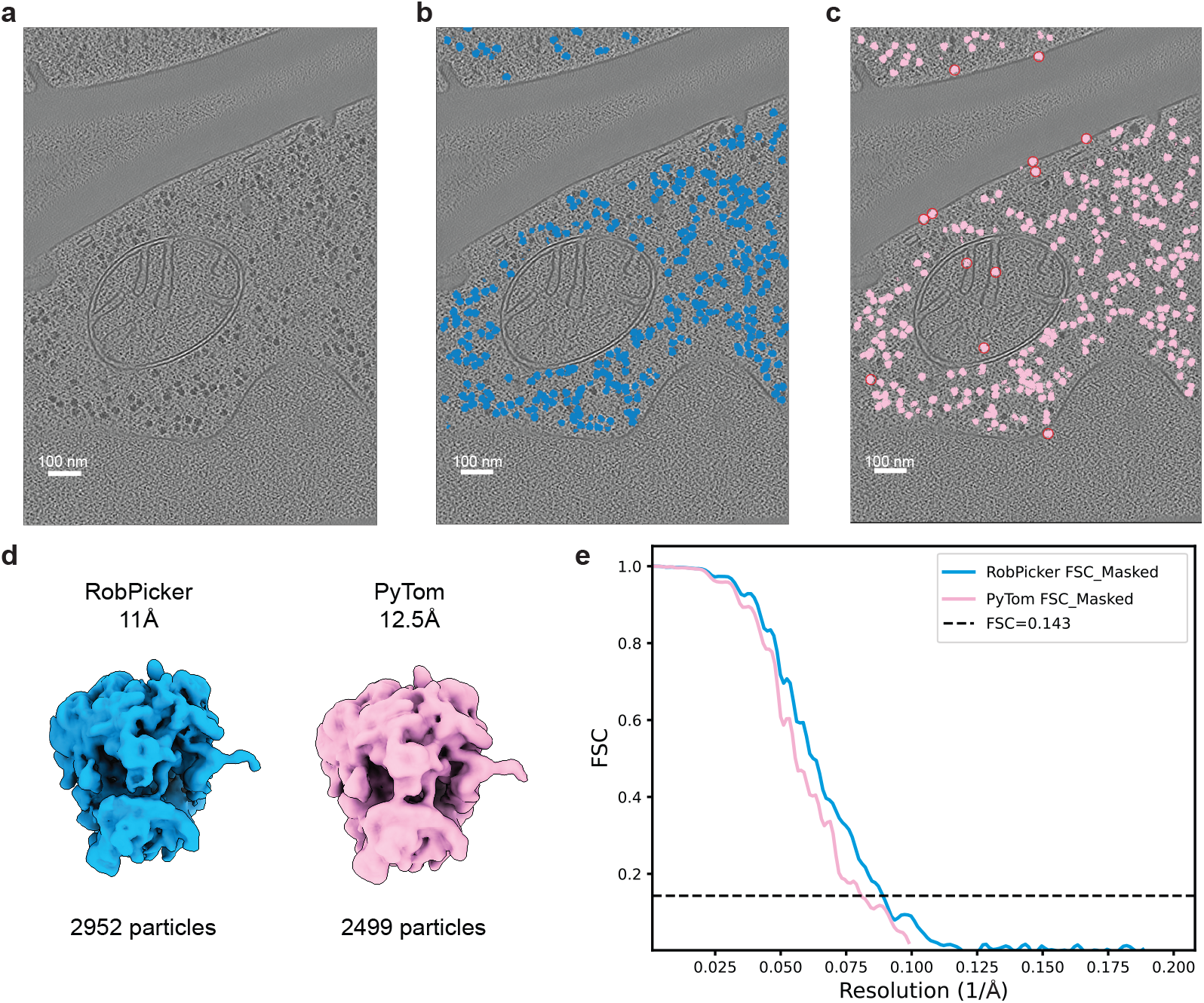
RobPicker provided supervision-free particle picks, compared with PyTom. **a**, Tomographic slice of a representative tomogram from yeast. **b**, Tomographic slice with ribosome picks by RobPicker highlighted. A total of 1,116 particles were found on this tomogram. **c**, The same slice with the top 1,116 candidates from PyTom template-matching highlighted. Typical false positive picks are marked in red circles. **d**, RELION refinement and classification as well as M refinement were used on both populations of particles to maximize the resolution. RobPicker produced a better reconstruction with more high-quality particles. **e**, FSC curves of the reconstructions shown in (d).

### RobPicker robustly picks particles of diverse sizes in noisy, imbalanced data

We conducted experiments on a benchmark dataset from the SHREC 2019 Cryo-ET Challenge, which consists of ten synthetic tomograms with ground truth labels for particles across 12 distinct species, covering a wide range of sizes [50]. These particles were categorized by size into four groups: large, medium, small, and tiny (Fig. 6). The dataset was divided into a training set, a validation set, and a test set (Methods). The label maps for the validation and test sets were created using the shape-based method, while the label maps for the training set were created based on two settings, one with the shape-based method and the other with the sphere-based method—to demonstrate the robustness of RobPicker facing data with different noise levels.

**Fig. 6:**
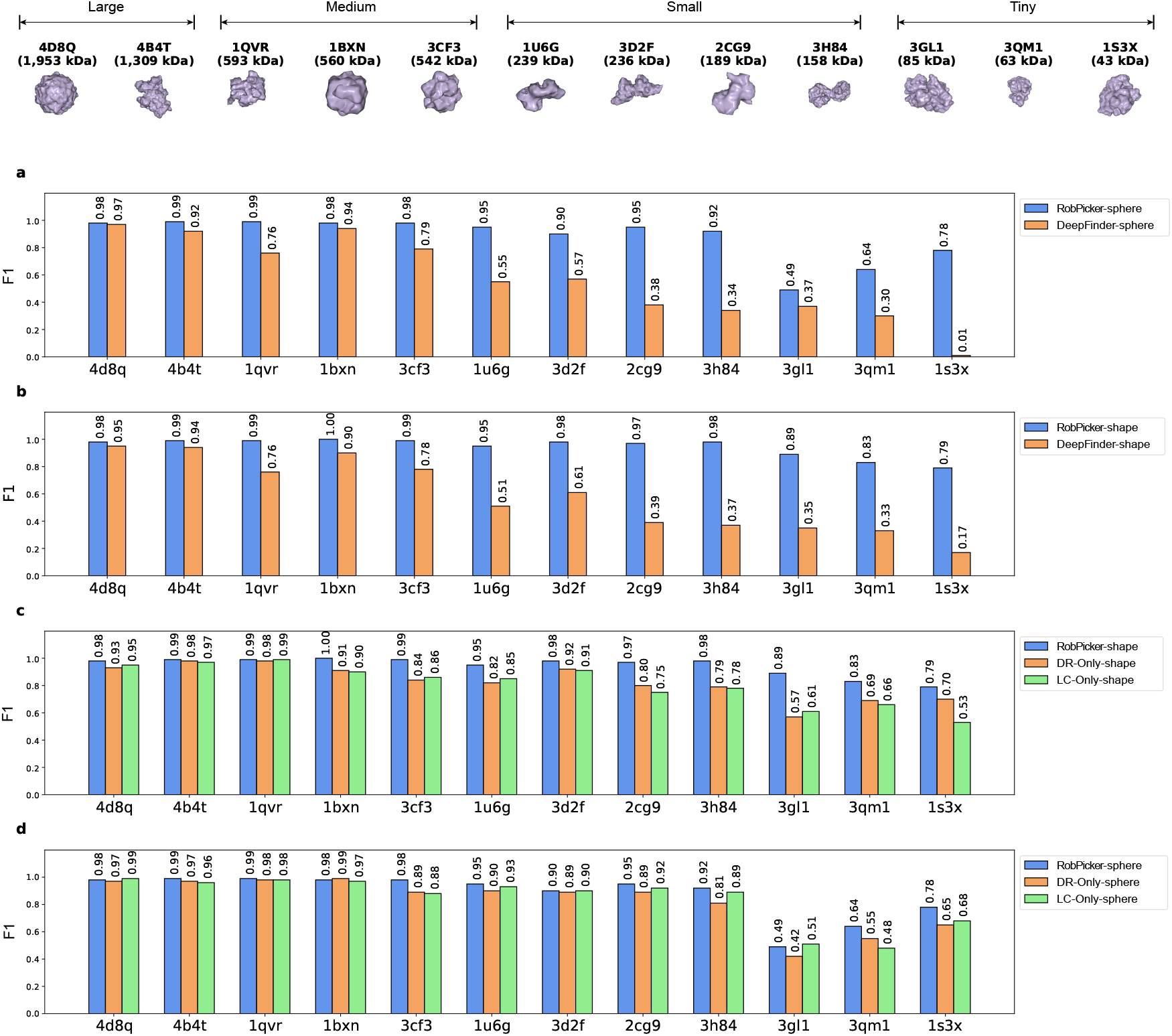
RobPicker significantly outperformed DeepFinder on the SHREC 2019 dataset. **a, b**, RobPicker outperformed DeepFinder across 12 distinct classes encompassing particles of different sizes. **a**, The training labels were sphere-based, where a sphere is used to approximate the occupancy of a particle. **b**, The training labels were shape-based, where particle contours were precisely used to annotate tomograms. The sphere-based labels were noisier than the shape-based labels but cheaper to generate. **c, d**, RobPicker integrated with both data reweighting network (DRN) and label correction network (LCN) outperformed the baselines of RobPicker with only DRN or LCN, in both shape- and sphere-label experiments.

We compared RobPicker’s performance with DeepFinder’s [20] in the benchmark. RobPicker consistently outperformed DeepFinder in terms of picking F1-scores across all particle size categories, including large, medium, small, and tiny in the sphere-based experiments (Fig. 6a). Notably, RobPicker demonstrated a substantial improvement over DeepFinder in detecting small and tiny particle species. For small species, such as 1u6g, 3d2f, 2cg9, and 3h84, RobPicker achieved F1 scores of 0.95, 0.90, 0.95, and 0.92, respectively, in contrast to DeepFinder’s scores of 0.55, 0.57, 0.38, and 0.34. This translates to relative improvements of 73%, 58%, 150%, and 171%, respectively. For tiny species like 3gl1, 3qm1, and 1s3x, RobPicker attained F1 scores of 0.49, 0.64, and 0.78, surpassing DeepFinder’s corresponding scores of 0.37, 0.30, and 0.01, with relative improvements of 32%, 113%, and 77-fold for these species. Notably, while DeepFinder struggled to identify 1s3x, achieving an F1-score of only 0.01, RobPicker excelled with a much higher score of 0.78. For medium-sized species such as 1qvr, 1bxn, and 3cf3, RobPicker achieved near-optimal F1-scores of 0.99, 0.98, and 0.98, respectively, outperforming DeepFinder’s corresponding scores of 0.76, 0.94, and 0.79, with improvements of 30%, 4%, and 24%. Similarly, for large particles like 4d8q and 4b4t, RobPicker attained F1-scores of 0.98 and 0.99, compared to DeepFinder’s 0.97 and 0.92.

RobPicker also outperformed DeepFinder in the shape-based experiments (Fig. 6b) where the training labels were cleaner than the sphere-based experiments. Moreover, RobPicker demonstrated a more pronounced improvement over DeepFinder for smaller particles compared to larger ones. Specifically, in the shape-based experiments, the average relative improvement over DeepFinder for tiny, small, medium, and large species is 224%, 115%, 23%, and 4%, respectively, revealing a decreasing trend. This can be attributed to the fact that smaller particles occupy fewer voxels (Fig. 1b), which leads to lesser influence on the optimization of the Dice loss function during model updates. Therefore, RobPicker’s data reweighting network is more effective on smaller particles than on larger ones, resulting in larger improvement for the smaller ones.

### Data reweighting and label correction networks both contribute to RobPicker’s robustness

To evaluate the effectiveness of the data reweighting (DR) network and label correction (LC) network, we tested two variants of RobPicker: DR-Only, which performs data reweighting without label correction, and LC-Only, which applies label correction without data reweighting. RobPicker with both DR an LC networks significantly outperformed the two variants (Fig. 6c,d). For picking particles of tiny species, RobPicker achieved F1 scores that were 56.1% and 45.9% (relative improvement) higher than DR-Only and LC-Only respectively on 3gl1, 20.3% and 25.8% higher on 3qm1, and 12.9% and 49.1% higher on 1s3x (Fig. 6c). Similarly, for small species, RobPicker outperformed DR-Only and LC-Only by 15.9% and 11.8% respectively on 1u6g, 6.5% and 7.7% on 3d2f, 21.3% and 29.3% on 2cg9, and 24.1% and 25.6% on 3h84. The improvement was also substantial for some medium species, with RobPicker achieving 9.9% and 11.1% higher scores on 1bxn, and 17.9% and 15.1% higher scores on 3cf3. These results highlight the importance of combining data reweighting and label correction to achieve robust performance in RobPicker.

The improvement of RobPicker over the two variants is more pronounced for smaller particles compared to larger ones. Specifically, the average relative improvement of RobPicker over DR-Only for tiny, small, medium, and large species is 29.8%, 16.9%, 9.2%, and 3.2%, respectively, showing a decreasing trend. Similarly, the average relative improvement of RobPicker over LC-Only follows a similar pattern, with the average relative improvement being 40.2%, 18.6%, 8.7%, and 2.6% for tiny, small, medium, and large species, also reflecting a decline as the particle size increases. These findings align with those in the previous section, underscoring the effectiveness of RobPicker in picking small and rare particles.

## Discussion

RobPicker’s improved robustness over DeepFinder and DeePiCt in identifying diverse macromolecule species can be attributed to several key factors inherent in its design. Unlike DeepFinder and DeePiCt, which rely on standard supervised learning approaches, RobPicker incorporates two crucial mechanisms—data reweighting and label correction—to address the challenges of species imbalance and noisy labels, which are not adequately handled by supervised learning methods.

One key component of RobPicker is its data reweighting network. In cryo-ET datasets, macromolecules often exhibit substantial species imbalance, with larger, more prominent species being overrepresented in the training data compared to smaller, rarer ones. Smaller, less abundant macromolecules (compared to ribosomes) can play pivotal roles in molecular pathways and other biomedical processes. For example, FAS has been found as the main enzyme cancer cells rely on to make the fats they need for growth and thus offers insight into anticancer therapy [51]. Accurately identifying these rare macromolecules allows researchers to reveal how cells respond to drugs, growth signals, or immune challenges. However, DeepFinder and DeePiCt, without reweighting mechanisms, are likely to learn biased representations, skewing their detection capabilities toward the dominant macromolecule species. This is especially evident in D5, where DeePiCt has a much lower performance on the small and rare species of FAS than on the abundant ribosomes. In contrast, RobPicker’s reweighting network automatically learned to elevate the weights of tomograms from underrepresented species, which increases their influence on training the segmentation network. Notably, the data weights are adjusted such that the model’s segmentation performance on the validation set is maximized, which makes it more suitable than the method of manually adjusting the weights of data samples as manual adjustment may not lead to optimal model performance. As a result, our data reweighting method leads to more balanced detection accuracy across both large and small macromolecule species, effectively addressing the bias that hampers traditional supervised learning methods.

Additionally, RobPicker’s label correction network further distinguishes it from DeepFinder and DeePiCt. Automatic labeling in cryo-ET data is often noisy due to the inherently low SNR of tomograms. Expert annotation is typically more accurate but requires substantial effort and time. Moreover, while the shape-based labeling method produce cleaner label maps, it is more time-consuming to generate than the sphere-based method. Therefore, human labor and data preprocessing time can be significantly reduced if noisy labels are effectively leveraged to train macromolecule segmentation networks. While DeepFinder and DeePiCt are prone to noisy labels, RobPicker corrects the noisy labels before training the segmentation network with them, significantly reducing the negative impact of labeling noise. This strategy leads to robust and improved detection of macromolecules, even with noisy training datasets.

Our results demonstrate that RobPicker can serve as an efficient, supervision-free entry point for cryo-ET structure determination pipelines. By eliminating the need for manual input or organism-specific hyper-parameter tuning, RobPicker streamlines particle detection while maintaining high specificity and generalizablity, enabling rapid generation of high-quality particle maps. In the yeast datasets, RobPicker outperformed the template-matching method PyTom [48], producing reconstructions of higher resolution and with fewer false positives. Importantly, its synergy with state-of-the-art refinement tools such as RELION [47, 52] and M [42] underscores how accurate particle picking can amplify downstream improvements in tilt-series alignment and map resolution. Together, these findings establish RobPicker not only as a robust particle picker but also as a practical tool that accelerates structural discovery in cryo-ET.

While RobPicker marks an improvement in macromolecule identification in cryo-ET, it has some limitations. A key challenge is its reliance on a small, clean validation dataset to guide the training of the label correction network. The scarcity of high-quality labels, often due to the manual effort required, may constrain the framework’s performance in settings with extremely limited well-annotated data. Additionally, RobPicker’s data reweighting network and label correction network, while effective and small (model sizes are detailed in Methods), introduce more memory requirement and computational complexity, as the iterative training of the networks can be resource-intensive, especially when using large-scale datasets. Furthermore, the current use of a 3D U-Net for macromolecule identification may not be ideal for cryo-ET data collected from different sources. Lastly, RobPicker primarily addresses noisy labels and data imbalance but does not explicitly account for variability in cryo-ET imaging conditions or sample preparation, limiting its generalizability to tomograms created in different settings.

Several future directions can be pursued to further enhance RobPicker’s performance and broaden its applicability in cryo-ET data analysis. One key area is reducing the reliance on clean validation datasets. Future efforts could explore semi-supervised or self-supervised learning techniques [53] for enabling RobPicker to better utilize noisy data. This would allow the framework to operate effectively in situations where obtaining high-quality labels is difficult and costly. Exploring alternative model architectures could also enhance RobPicker’s capabilities. Vision Transformer-based models with attention mechanisms [54], which have shown great success in image classification and segmentation [55], may offer superior performance over the current 3D U-Net when dealing with highly complex and variable cryo-ET data. Such architectures could improve the detection of challenging macromolecule species, particularly those with subtle features or lower SNR. Additionally, integrating methods to address variability in cryo-ET data, such as imaging noise or artifacts from sample preparation, could further increase RobPicker’s robustness. Developing preprocessing techniques that standardize tomograms before detection could mitigate these issues and enhance the model’s generalizability across diverse datasets. Finally, extending RobPicker’s framework to other imaging modalities, such as cryo-electron microscopy (cryo-EM) or super-resolution microscopy, could broaden its application scope.

## Methods

### The RobPicker framework

RobPicker is formulated as a meta learning problem, with a bi-level optimization (BLO) formulation. A BLO problem has two levels of optimization problems: a lower level and a upper level. In RobPicker, the training of the macromolecule segmentation network is at the lower level, while the training of the DR network and LC network is at the upper level. The two levels are nested and mutually dependent on each other: the optimal parameters of the segmentation network from the lower level are used to define the objective function at the upper level; the non-optimal parameters of the DR and LC networks from the upper level are used to define the objective function at the lower level. Due to this mutual dependency, the model parameters of the two levels are iteratively updated, with the detailed process described below.

Given a training dataset 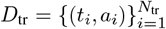, where *t*_*i*_ represents an input tomogram, *a*_*i*_ denotes the corresponding macromolecule label, and *N*_tr_ is the number of training samples, we input *t*_*i*_ into the macromolecule segmentation network *f* (*t*_*i*_; *I*) with *I* representing the network’s weight parameters, which outputs a prediction. Simultaneously, *a*_*i*_ is fed into the LC network *g*(*a*_*i*_; *C*), where *C* represents the weight parameters for this network, producing a corrected label. A loss function *ℓ*(*f* (*t*_*i*_; *I*), *g*(*a*_*i*_; *C*)), which is the Dice loss [56], is computed to quantify the discrepancy between the detection result and the corrected label. The loss value is fed into the DR network to output a scalar weight *h*(*ℓ*(*f* (*t*_*i*_; *I*), *g*(*a*_*i*_; *C*)); *R*), where the DR network is parameterized by *R*. The loss is then multiplied by this weight. Let *L*(*D*_tr_, *I, C, R*) represent the sum of the reweighted losses over the entire training dataset:

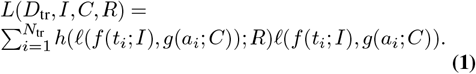

We train the segmentation network by updating its parameters *I* to minimize this loss *L*(*D*_tr_, *I, C, R*) while keeping the parameters of the LC and DR networks fixed. This leads to the following optimization problem:

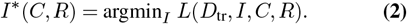

Here, *I*\*(*C, R*) indicates that the optimal trained parameter *I*\* is a function of the parameters of the LC and DR network *C* and *R*, as *I*\* is determined by the loss function, which itself depends on *C* and *R*. Note that *C* and *R* are not optimized in this step, as doing so could result in a trivial solution where the weight *h*(*ℓ*(*f* (*t*_*i*_; *I*), *g*(*a*_*i*_; *C*)); *R*) becomes zero for every *i*.

Next, we evaluate the trained segmentation network *I*\*(*C, R*) on a clean validation dataset 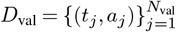, where *N*_val_ represents the number of validation examples. For each validation tomogram *t*_*j*_, the segmentation network is used to detect macromolecules, and the resulting prediction *f* (*t*_*j*_; *I*\*(*C, R*)) is compared to the corresponding ground truth label *a*_*j*_, yielding a loss *ℓ*(*f* (*t*_*j*_; *I*\*(*C, R*)), *a*_*j*_). Let *L*(*D*_val_, *I*\*(*C, R*)) represent the total loss over the entire validation dataset:

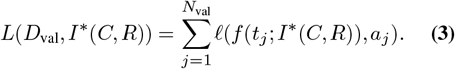

We optimize the parameters of the CL and DR networks *C* and *R* by minimizing the validation loss, which leads to solving the following optimization problem:

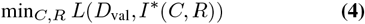

Combining Equations 2 and 4, we obtain the following bilevel optimization problem:

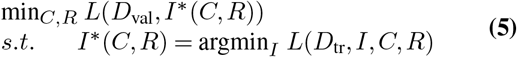

The two levels of optimization problems are interdependent. The solution from the lower level, *I*\*(*C, R*), is used to define the loss function at the upper level, while the optimization variables *C* and *R* from the upper level are also involved in defining the loss function at the lower level.

### Optimization algorithm

We address the optimization problem in Equation 5 using a hypergradient-based approach. Specifically, we approximate the optimal solution *I*\*(*C, R*) through a single-step gradient descent update as follows:

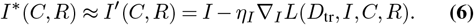

where *η*_*I*_ denotes the learning rate.

We substitute the approximation *I*\*(*C, R*) ≈*I*′(*C, R*) into the upper-level loss function, yielding an approximate loss *L*(*D*_val_, *I*′(*C, R*)). We then update *C* and *R* using gradient descent with respect to the approximate loss:

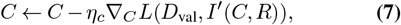

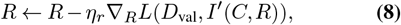

where *η*_*c*_ and *η*_*r*_ denote the respective learning rates. The gradient ∇_*C*_*L*(*D*_val_, *I*′(*C, R*)) can be computed using the chain rule:

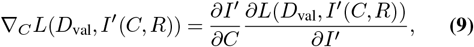

Where

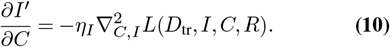

Similarly, the gradient ∇_*R*_*L*(*D*_val_, *I*′(*C, R*)) is computed as:

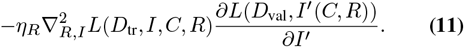

The updates in Equations 6, 7, and 8 are performed iteratively until convergence. The steps of the algorithm are outlined in Algorithm 1.

In Equation 11, computing the matrix 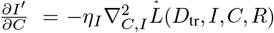 and performing multiplication with the vector 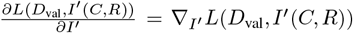 incur noticeable computational cost. To mitigate this, we adopted the finite difference approximation method [57]. This method is a numerical technique for estimating derivatives when their analytical forms are either complex or infeasible to compute. By evaluating the function at perturbed points, finite difference approximation provides an efficient way to approximate derivatives. Let *I*^+^ and *I*^−^ denote *I* + *ϵ*∇_*I*_*′ L*(*D*_val_, *I*′(*C, R*)) and *I* − *ϵ*∇_*I*_*′ L*(*D*_val_, *I*′(*C, R*)), respectively, where *ϵ* is a small scalar. Using this approach, the matrix-vector multiplication can be approximated as:

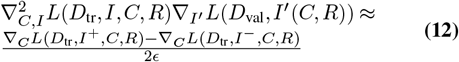

Similarly, 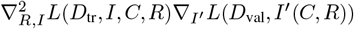 can be approximated as:

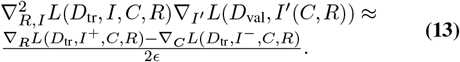

This bi-level optimization algorithm has been implemented in our Python library Betty [58], based on which RobPicker was implemented.

### Model architecture

We used PyTorch [59] to implement the macromolecule segmentation network. For the experiments in comparison with DeepFinder, we used the 3D U-Net [24] following DeepFinder [20]. The input of the macromolecule segmentation network is a 3D tensor (e.g., *h* × *w* × *d*), representing a 3D tomogram. The output of the macromolecule segmentation network is a 4D tensor (*h* × *w* × *d* × *c*) with *c* channels, where *c* matches the number of classes (number of species plus one for the background). The model has four down-blocks, where each down-block has four 3D convolutional layers (kernel size: 3×3×3, stride: 1×1×1, padding: 1×1×1). Each convolutional layer is followed by a 3D batch normalization layer [60] and a ReLU activation function [61]. There is a 3D max-pooling (kernel size: 2, stride: 2, padding: 1, dilation: 1) between the second and the third convolutional layer in each down-block. The model essentially predicts the probabilities of each voxel belonging to each class. If we perform the argmax operation along the channel dimension, we can get the predicted label of each voxel belonging to one of the classes (see post-processing below). The down-blocks are followed by a bottleneck block with two 3D convolutional layers, which is followed by four up-blocks. Each upblock has a transposed 3D convolutional layer (kernel size: 2×2×2, stride: 2×2×2) and two 3D convolutional layers (kernel size: 3×3×3, stride: 1×1×1, padding: 1×1×1). The final layer of this segmentation network is 3D convolutional layer (kernel size: 1×1×1, stride: 1×1×1). For the experiment on D5 where RobPicker was compared with DeePiCt, we used the same 3D U-Net as DeePiCt [21], which has a similar architecture as DeepFinder and it has 41 million parameters.

#### Algorithm 1

Bi-level optimization for RobPicker

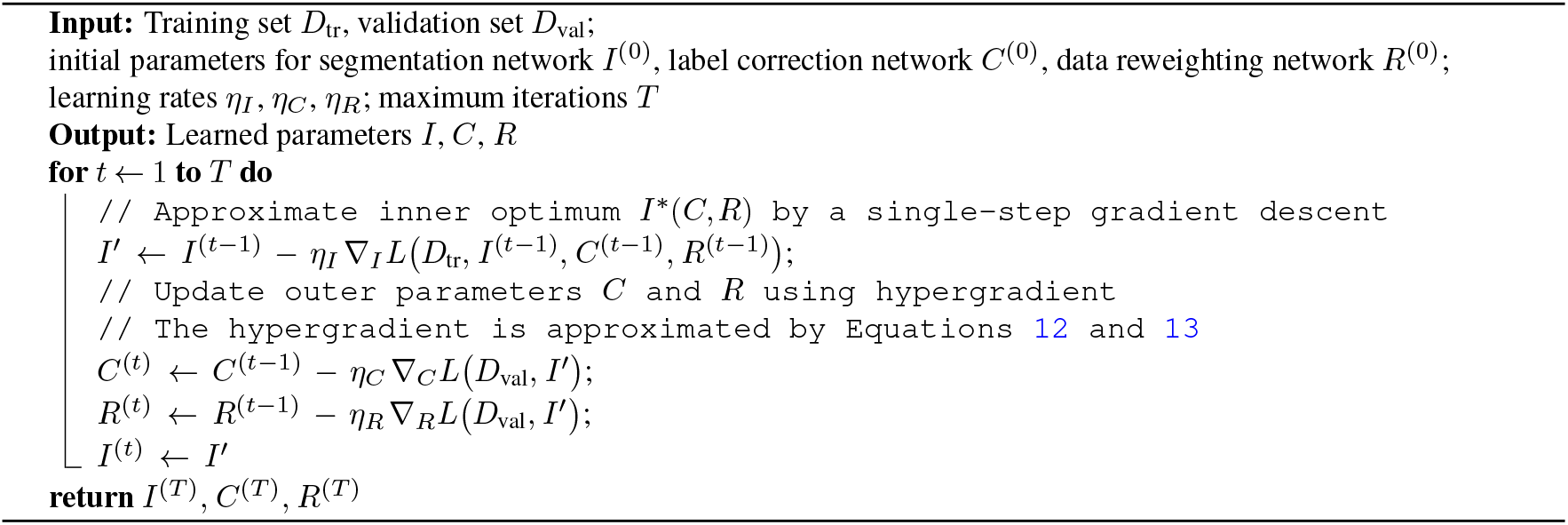

The data reweighting network has a linear layer that expands the input dimension of 1 (i.e., the loss value) to a hidden dimension of 500, followed by a ReLU activation [61], and then a linear layer that shrinks the hidden dimension of 500 to the output dimension of 1, followed by a sigmoid function that converts the output to a weight between 0 and 1. The data reweighting network has only 1.5 thousand parameters. The label correction network takes as input the noisy label, denoted by **L** (4D tensor with *c* channels, where the channel dimension is one-hot), and the input tomogram (3D tensor). The label correction network starts with a sub-network that processes the input tomogram and outputs a feature map of the tomogram, denoted by **M**. The sub-network shares the same architecture and parameters as the macromolecule segmentation network for efficiency consideration. A 3D convolution layer maps the input channel size of **M** to the output channel size of the number of classes (with kernel size being 1, padding 1, and stride 1), followed by a softmax operation. The result of the softmax is denoted by **S**. Then, **S** is concatenated with the noisy label. The result feature map has a channel size of two times the number of classes. Then, a 3D convolution layer (with kernel size being 1, padding 1, and stride 1) shrink the channel size to the number of classes, followed by a ReLU activation, another 3D convolution layer (with kernel size being 1, padding 1, and stride 1), and a sigmoid operation. The result, denoted by *α*, acts as a mixing coefficient of the noisy label **S** and the softmax result **S** to compute the corrected label **C**:

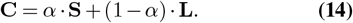

Due to the sharing of parameters between the label correction network and the macromolecule segmentation network, the label correction network only has around 1.3 thousand parameters. The small amount of parameters of the label correction and data reweighting networks has significantly reduced the computational complexity of the bi-level optimization in RobPicker.

### Data preprocessing

#### Data structure

A tomogram is represented as a 3D tensor with shape *h* × *w* × *d*, while its label map is represented as a 4D tensor with shape *h* × *w* × *d* × *c*, where *c* is the number of classes for segmentation. For a label map denoted by *M*, and for a specific 3D coordinate (*x, y, z*), we have a ground-truth class, denoted by *e* ∈ {1, 2,…, *c*}. Then, the label map is constructed so that *M*_*x*,*y*,*z*,*e*_ = 1 and *M*_*x*,*y*,*z*,*j*_ = 0 for *j* ≠ *e*.

#### Tomogram patching

In the training phase of deep learning models for tomographic data of large sizes (e.g., 500 × 500 × 500), GPU memory capacity often becomes a bottleneck, often causing out of memory error. To circumvent this limitation, we divided a whole tomogram into distinct 3D subtomograms, or “patches”. Following DeepFinder [20], we used a patch size of 64 × 64 × 64. We also applied random rotation and random shift to the input for data augmentation during training. The labels were extracted using the same patch size accordingly. Therefore, the segmentation network only needs to take a 3D subtomogram as input and predict the label on the subtomogram. The advantage of this strategy lies in its memory efficiency—only the batch currently under scrutiny is loaded into the GPU memory. This ensures that our methodology remains adaptable across a spectrum of GPUs. During inference, the test tomogram is also divided into subtomogram to input to the segmentation network. However, the prediction on the boundary of the subtomograms can be less accurate due to the lack of sufficient tomographic context. Therefore, we extrated overlapping subtomograms (with an overlap of 20 voxels) and averaged the overlapping regions of the predicted segmentation maps.

#### Resampling

To make the particle classes (i.e., macromolecule species) distributed balanced in the validation data, we used resampling when extracting the subtomograms. Specifically, we first uniformly sample a particle class from {1,2,…, c} and then sample a particle from the chosen particle class. Then, the subtomogram with the sampled particle is extracted for training.

### Data post-processing

Similar to the methodology adopted by DeepFinder [20], RobPicker’s post-processing operation is a multi-step procedure involving classification and clustering. Applying the macromolecule segmentation network on the input tomogram results in a segmentation map, which is denoted by *M*. Then, the first step is classification of every voxel within the tomogram—ensuring each voxel is attributed to a specific macromolecule species—by selecting the class with the highest predicted probability. Mathematically, we compute *C*_*x*,*y*,*z*_ = argmax_*i*_ *M*_*x*,*y*,*z*,*i*_ to obtain a 3D tensor *C*, which effectively becomes the label map predicted by the model.

Building on the voxel classification, RobPicker’s subsequent focus is clustering. In this phase, a robust clustering algorithm, called DBSCAN [62], groups the classified voxels into distinct units that represent individual particles. For the configuration of the DBSCAN algorithm, we set the radius of a neighborhood to 1 voxel and minimum number of voxels in a cluster to 5. After clustering, the exact location of each particle is revealed within the tomogram volume by calculating the gravity center of the voxel clusters. For the 3D visualization of the particles, we use IMOD [63], Chimera [64], and ChimeraX [65].

### Datasets

D1 was obtained from DeepFinder [20]. It encapsulates 57 tomograms of *C. reinhardtii* cells. Experts manually annotated the 3D coordinates of 8,792 mb-ribos using a combination of template matching, subtomogram classification, and visual inspection. Ct-ribos were annotated by semiautomatic tools without expert supervision.

D2 was generated to study co-translating 80S ribosomes in yeast cells [46]. It includes 6 tomograms from yeast lamellae, where PyTom [48] template-matching and manual expert annotations were used to annotate 80S ribosomes. Tomograms were collected at the pixel size of 2.63 Å and reconstructed at the pixel size of 10 Å in Warp [66]. All tomograms were denoised in IsoNet [29] to compensate for the missing wedge effects.

For D3, we collected 28 tomograms from *E. coli* cells. These tomograms were collected at pixel size of 1.66 Å and reconstructed and binned by 6 voxels at 9.98 Å using Warp and were not denoised any further. The annotation for this dataset was done through template-matching and classification in RELION [47] to separate the 70S and 50S classes.

For D4, we collected 5 tomograms using the following procedure. Yeast cells in the logarithmic growth phase were vitrified and prepared for cryo-ET using cryo-FIB milling [67]. A ribosome particle picker was trained on this dataset and subsequently applied to eight additional tomograms of yeast (also prepared using cryo-FIB) without any supervision, yielding ∼23,000 ribosome coordinates. These particles were subjected to refinement in RELION 3 [47]. The workflow included an initial refinement of all particles, followed by 3D classification. Particles belonging to the best-resolved class (∼5,500 particles) were selected for subsequent refinements, resulting in a good and representative yeast ribosome map (resolution ∼19 Å) shown in Fig. 4b.

The *M. pneumoniae* dataset that was used to demonstrate the pipeline in Fig. 4c was obtained from M [42]. The same RobPicker trained on D4 was used to pick ∼5,300 particles from 14 tomograms of *M. pneumoniae*. These picks were subjected to initial refinement and classification in RELION 3 [47]. From these, ∼2,000 particles belonging to the highest-quality class were selected for multi-particle refinement in M [42]. This refinement, coupled with tilt-series alignment fine-tuning, improved the ribosome map resolution from ∼15 Å to 7 Å (Fig. 4c). All reported resolutions were estimated using the gold-standard Fourier shell correlation (FSC) at the 0.143 criterion.

D5 was obtained from DeePiCt [21]. It includes 10 VPP tomograms of *S. pombe* cells. An iterative workflow was used to localize ribosome and FAS in 4×-binned tomograms (13.48 Å voxel size). Manually curated template matching was performed for ribosomes, and non-exhaustive manual picking was performed for FAS (step 1). The resulting annotations were used to train the 3D CNNs of DeePiCt (step 2). For ribosomes, step 2 was repeated three times (always trained on combined predictions of step 1 and the preceding round). Cumulative predictions were manually revised in tom_chooser in PyTom (for ribosomes) and in EMAN2 [68] (for FAS). This comprehensive pipeline ensured the labels were high-quality.

The SHREC 2019 challenge [50] utilized 12 Protein Data Bank (PDB) identifiers (1bxn, 1qvr, 1s3x, 1u6g, 2cg9, 3cf3, 3d2f, 3gl1, 3h84, 3qm1, 4b4t, and 4d8q) to generate tomogram density maps. Then, the density maps were subsequently transformed into a tilt series of projection images, which were then deliberately degraded with noise and contrast adjustments. The reconstructed tomograms from tilt series have a dimension of 512 × 512 × 512 with a 1 nm voxel resolution. Each tomogram contains approximately 200 particles per species on average. Nine tomograms were provided for training, while the tenth tomogram was reserved for testing. For RobPicker, we partitioned the nine training tomograms into two subsets, assigning eight tomograms for training and one for validation. These subsets were employed in Stage I and Stage II—respectively—of the training process shown in Fig. 1c.

### Experimental Settings

#### Loss function

We used the Dice loss function following DeepFinder [20] and DeePiCt [21]. The Dice loss function is computed for each channel (i.e., class) and the final loss value is the averaged loss values of all the channels. Specifically, for a ground truth label map *A* (4D tensor of shape *h* × *w* × *d* × *c*) and a predicted label map *B*, whose values are in the interval [0, 1], the Dice loss is computed as:

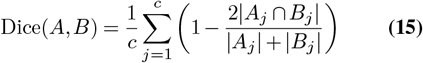

where *j* is the index for the last dimension, and the intersection 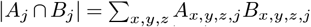, and the union 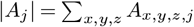 and 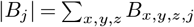.

#### Evaluation metrics

We used the picking F1 score to assess particle picking performance. The F1-score is the harmonic mean of precision and recall:

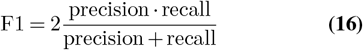

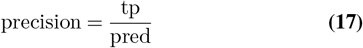

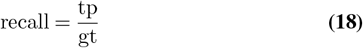

where tp, pred, gt denote the numbers of true positives, predicted particles, and ground truth particles, respectively. A predicted particle was considered to be a true positive if the centroid of the predicted particle was located within the boundary of a ground truth particle with the same class label. For the evaluation of model performance on D5 from DeePiCt [21], we followed DeePiCt’s evaluation metric to define the true positives as those predicted particles whose coordinates overlap with a ground truth particle within a tolerance radius (10 voxels, 135 Å).

#### Hyperparameters and Optimization

The stochastic gradient descent (SGD) algorithm [69, 70] was used to update model parameters. SGD encompasses a learning rate of `lr=0.0001`, momentum parameterized at `momentum=0.9`, and a weight decay set to `weight_decay=0.0005`. Efficient training was facilitated with batches of size `batch_size=25`. The model was subjected to 10,000 training iterations, as captured by `train_iters=10000`, with validation step performed at every 500 iterations, `valid_iters=500`. The metalearning facets of our architecture utilized parameters such as `meta_alpha=0.0`, and `lamda=1.0`, among other configurations which are available in the documentation of our code. For a dataset with 10 tomograms, the training process typically takes a few hours on an Nvidia Tesla A100 (80 GB) GPU.

## Data availability

D1 can be found in the Electron Microscopy Data Bank (EMDB) under accession number EMD-3967. D2 is available at EMPIAR-12534. D3 and D4 were generated by ourselves and pending release. D5 is available at EMPIAR-10988. The *M. pneumoniae* tomogram dataset is available at EMPIAR-10499. The dataset of the SHREC 2019 challenge can be accessed at the following URL: http://www2.projects.science.uu.nl/shrec/cryo-et/2019/.

## Code availability

The source code of RobPicker is available at https://github.com/RobPicker/CryoET-RobPicker

---

**Extended Data Figure 1:**
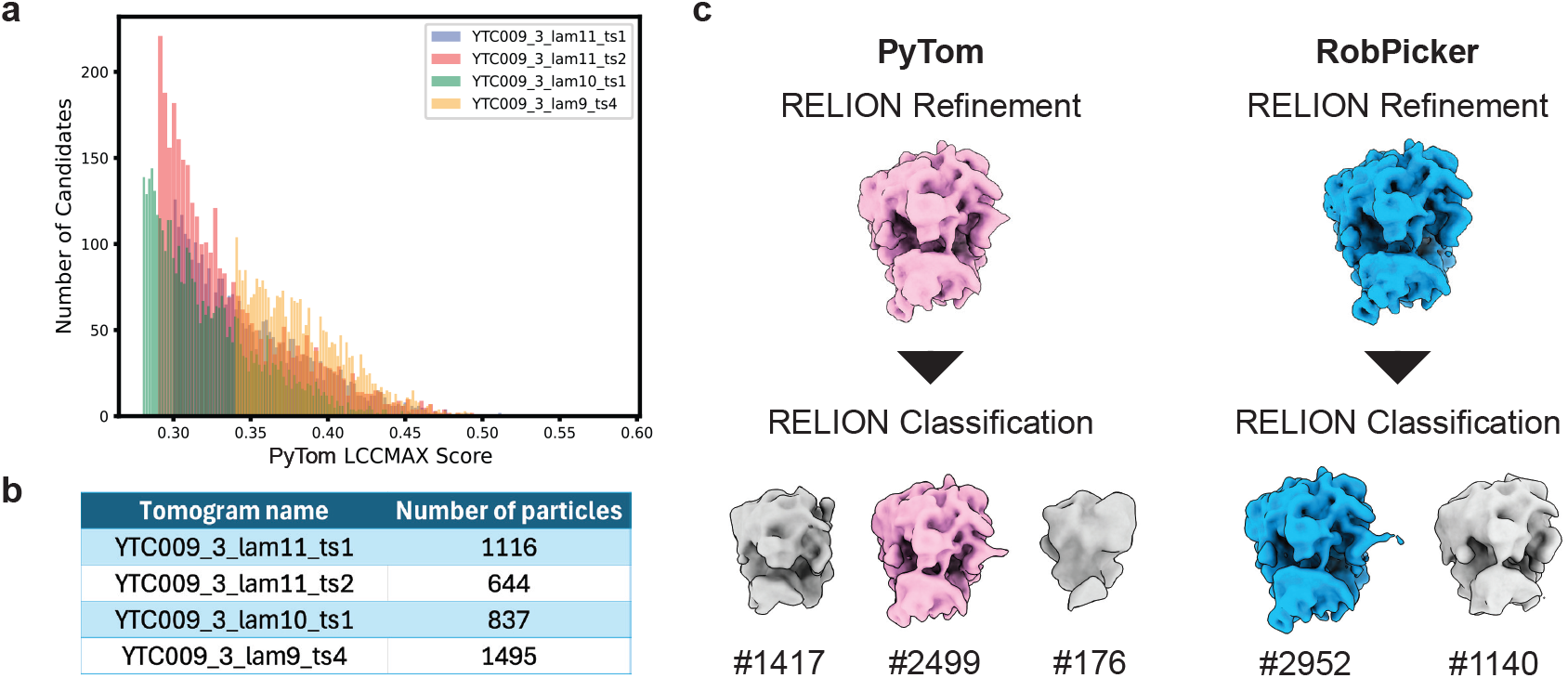
RobPicker is supervision-free in structure solving for yeast ribosomes in D2. **a**, Histogram of the LCC scores from PyTom for the top 10,000 particles does not show the desired distribution with peaks for automatic thresholding. **b**, Number of ribosomes for each tomogram picked by RobPicker which does not need any manual input. **c**, Top PyTom particles from the 4 tomograms are extracted and compared before and after classification. The best classes then are refined in RELION and M to produce final structures shown in Fig. 5d.

## Notes

### Competing Interest Statement

The authors have declared no competing interest.

